# A novel methodology for defining stromal expression of atypical chemokine receptors in vivo

**DOI:** 10.1101/748673

**Authors:** Christopher A.H. Hansell, Samantha Love, Marieke Pingen, Gillian J Wilson, Megan MacLeod, Gerard J. Graham

## Abstract

Analysis of chemokine receptor, and atypical chemokine receptor, expression is frequently hampered by the lack of availability of high-quality antibodies and the species-specificity of those that are available. We have previously described methodology utilising Alexa-Fluor labelled chemokine ligands as versatile reagents to detect receptor expression. Previously this has been limited to haematopoietic cells and methodology for assessing expression of receptors on stromal cells has been lacking. Amongst chemokine receptors the ones most frequently expressed on stromal cells belong to the atypical chemokine receptor subfamily. These receptors do not signal in the classic sense in response to ligand but scavenge their ligands and degrade them and thus sculpt in vivo chemokine gradients. Here we demonstrate the ability to use either intratracheal, or intravenous, Alexa-Fluor labelled chemokine administration to detect stromal cell populations expressing the atypical chemokine receptor ACKR2. Using this methodology we demonstrate, for the first time, expression of ACKR2 on blood endothelial cells. This observation sets the lung aside from other tissues in which ACKR2 is exclusively expressed on lymphatic endothelial cells. In summary therefore we described a novel method for the in situ labelling of atypical chemokine receptor expressing cells appropriate for subsequent flow cytometric analysis. We propose that this methodology will work in a range of species and for a range of receptors and therefore will have significant versatility

## Introduction

In vivo leukocyte migration is regulated, in the main, by proteins belonging to the chemokine family of chemotactic cytokines(1, 2). This family is defined on the basis of a conserved cysteine motif in the mature sequence of its members and is divided into CC, CXC, XC and CX3C subfamilies according to the specific configuration of this motif. The chemokine family arose early in vertebrate evolution(3) (pre-vertebrate species do not have chemokines) and the primordial chemokine was almost certainly CXCL12 which plays essential roles in stem cell migration during embryogenesis(4–9). From this one chemokine and its receptor CXCR4, through gene duplication, the family has expanded to the point at which mammals have approximately 45 chemokines, and 18 signalling chemokine receptors, which together orchestrate in vivo homeostatic and inflammatory leukocyte migration. Chemokine regulation of cellular migration is extremely complex and, particularly in the case of inflammation(10), poorly understood which has contributed to ongoing problems in therapeutically targeting inflammatory chemokine receptors in immune and inflammatory diseases(11).

In addition to signalling chemokine receptors which belong to the G-protein coupled receptor family(12), chemokines also bind to a subfamily of atypical chemokine receptors (ACKRs) which are generally stromally-expressed and which fine-tune in vivo chemokine activity by scavenging chemokines and therefore regulating chemokine availability(13–15). There are currently 4 members of the atypical chemokine receptor family. ACKR1 (formerly known as DARC); ACKR2 (formerly known as D6); ACKR3 (formerly known as CXCR7) and ACKR4 (formerly known as CCX-CKR). With the exception of ACKR1, these receptors exhibit spontaneous internalisation and recycling activity and scavenge chemokines from the environment and target them for lysosomal degradation. ACKR3 carries out this role in some essential developmental contexts and is strongly evolutionarily conserved(5, 6, 16, 17). ACKR4 scavenges chemokines within the lymph node to generate intra-lymph node gradients and facilitate dendritic cell migration from the sub-capsular sinus into the T cell zone of the lymph node(18). We have had a particular interest in ACKR2 which is the prototypic member of the atypical chemokine receptor family(19). This receptor binds, internalises and degrades all inflammatory chemokines belonging to the CC-chemokine subfamily and thus plays an essential role in the resolution of the inflammatory response(20–24). This has implications for tumorigenesis(25–28) as well as for branching morphogenesis in a number of developmental contexts(29, 30). ACKR2 is predominantly expressed on lymphatic endothelial cells(31, 32) and placental trophoblasts(23, 33, 34) with expression also being detected on subsets of splenic B cells(35). The atypical chemokine receptors are therefore central regulators of chemokine activity in vivo.

One of the challenges in studying chemokine receptors, including atypical chemokine receptors, is access to high quality antibodies. Whilst antibodies to some typical and atypical receptors are available, it has proven extremely difficult to raise useful antibodies to others. For example, there are no high-quality antibodies available for detection of murine ACKR2. A further issue is that even for receptors for which antibodies are available, these antibodies are invariably species-specific and it is therefore difficult to carry out analyses in either non-murine or non-human species. This is particularly important for highly conserved receptors such as ACKR3. The availability of a generic methodology to allow detection of atypical chemokine receptors (and amenable to detection of typical receptors), which would be usable in vivo and applicable to numerous species, would therefore represent a significant technological development.

Here we describe, a novel methodology for the detailed analysis of the phenotypes of ACKR2-expressing cells in lung stroma. The technology uses ACKR2-dependent internalisation of a fluorescently tagged ligand to identify receptor expressing cells and we demonstrate the utility of comparing intra-tracheal and intravenous administration in defining, and discriminating between, discrete receptor expressing stromal cell types. We propose that this methodology will be broadly applicable to a range of species and experimental analyses.

## Materials and Methods

### Mice

Animal experiments were performed using co-housed mice in ventilated cages in a barrier facility that conformed to the animal care and welfare protocols approved by the University of Glasgow under the revised Animal (Scientific Procedures) Act 1986 and the European Union Directive 2010/63/EU. Ackr2-deficient mice (Jamieson *et al*., 2005) were bred in-house (C57BL/6 background); wild type (WT) C57BL6/J mice were from Charles River Research Models and Services. Prox-1 reporter mice were obtained from Jackson laboratories. All experimental mice were sex and age matched.

### qPCR

RNA was extracted using RNeasy columns with DNase treatment (Qiagen), and the amount of RNA was quantified on a Nanodrop 1000 Spectrophotometer (Thermo Fisher Scientific). cDNA was synthesized using High-capacity RNA-to-cDNA kit by Applied Biosystems (Thermofisher). For all qPCRs, a final concentration of 0.2-mM primers was used for each PCR set up using PerfeCTa SYBR Green FastMix and ROX qPCR Master Mix (Quanta BioSciences). qPCRs were performed on a Prism 7900HT Fast Real-Time PCR System (Applied Biosystems). The thermal cycles for qPCR of TBP and ACKR2 were 95°C (3 min) for one cycle and 95°C (3 s) and 60°C (30 s) for 40 cycles. Relative expression was calculated using serial dilutions of cDNA standards. Primer sequences designed for qPCR and for producing cDNA standards were designed using Primer3 software (http://frodo.wi.mit.edu/cgi-bin/primer3/primer3_www.cgi). The following primers were used: mouse ACKR2, 5’-TTCTCCCACTGCTGCTTCAC-3’, 5’-TGCCATCTCAACATCACAGA-3’; mouse TBP primer: 5’-AAGGGAGAATCATGGACCAG-3’, 5’-CCGTAAGGCATCATTGGACT-3’.

### In-situ hybridisation

Mice were culled using increasing concentration of CO_2_. Lungs were placed in 10% neutral buffered formalin at room temperature for 24-36 hours before they were processed by dehydration using rising concentrations of ethanol, xylene stabilisation and paraffin embedding (Shandon citadel 1000, Thermo Shandon). Tissue was then sectioned onto Superfrost plus slides (VWR) at 6μm using a Microtome (Shandon Finesse 325 Microtome, Thermo). All slides for analysis were processed together. Slides were baked at 60°C for 1 hr before pre-treatment. Slides were deparaffinised with xylene (5 mins × 2) and dehydrated with ethanol (1 min× 2). In situ hybridisation was performed using the RNAscope® 2.5 HD Reagent Kit-RED from Advanced Cell Diagnostics (cat. no. 322350) and according to the manufacturer’s instructions. Briefly tissues were incubated with Hydrogen peroxide for 10 mins at RT. The slides were boiled in antigen retrieval buffer for 15 mins. Slides were treated with ‘protease plus’ for 30 mins at 40°C. Slides were then hybridised using the RNAScope 2.5 Red Manual Assay (Advanced cell diagnostics) according to manufacturer’s instructions using the Mm-ACKR2 probe (NM_021609.4). Slides were mounted in DPX (Sigma Aldrich) and imaged on an EVOS M7000 microscope (Thermofisher).

### Intratracheal and Intravenous Chemokine administration

To administer fluorescent chemokine intratracheally mice were euthanised using an appropriate schedule 1 method or CO_2_ exposure. The mice were then carefully dissected to remove the ribcage and expose the intact lungs and trachea *in situ*. Using a pair of surgical scissors, a small incision was made at the top of the exposed trachea toward the base of the jaw. A 2µg/ml solution of Alexa 647™ labelled CCL22 (Almac; Alexa-CCL22) dissolved in RPMI/25mM HEPES was prepared in a polypropolene tube and preserved from light at room temperature until needed. Once the dissection was complete a syringe with a 19G needle and loaded with 400µl of the Alexa-CCL22 solution was then inserted into the exposed trachea via the incision. The needle should be tight within the trachea and care should be taken not to pierce the trachea further down. The lungs were then inflated with the chemokine solution and the trachea carefully tied off with surgical thread to prevent the leakage of chemokine solution as the syringe is removed. The intact inflated lungs were then removed and placed into a falcon tube containing enough RPMI to cover the intact inflated lungs. The lungs were then incubated in a water bath for 1hour at 37°C. Following this time the lungs were removed from the waterbath and the surgical thread cut to allow draining of the remaining chemokine solution. The lungs were then digested for a single cell suspension as per the protocol.

### Flow cytometry

Lungs were removed and finely chopped with scissors, then incubated in digestion mix (1.6 mg/ml Dispase (Roche),0.2 mg/ml Collagenase P (Roche) and 0.1mg/ml DNase I (Invitrogen)) in HBSS on a gentle shake at 37°C for 40 mins. Lungs were passed through a 40µm mesh (Greiner Bio-One), RBCs were lysed in 1 ml RBC lysis buffer (eBioscience) for 1 min and then 10 mls FACs buffer (PBS, 1%FCS, 0.02% sodium azide, 5mM EDTA) to quench the reaction. The resulting single cell suspension was preincubated with Fc block (BD Biosciences) in FACS buffer (PBS, 1% FCS, and 5mM EDTA) and labeled with fluorescent anti-mouse Abs, including: CD45 (30-F11), Gp38 (8.1.1), CD140a (APA5), EpCAM-1 (G8.8), CD31 (390) and CD49f (GoH3) each labelled with various fluorochromes (Biolegend) CD166 (eBioALC48) (Ebioscience). Dead cells were labeled using Fixable viability dye efluor 506 or efluor 780 (ebioscience). Cells were stained for 20 minutes on ice before washing with FACS buffer. Cells were either analysed by flow cytometry immediately or fixed for 20 minutes with 2% methanol-free paraformaldehyde (Invitrogen) before washing and placing at 4°C in the dark until ready to be analysed typically 12-24 hours later. Cells were analyzed using LSR2 flow cytometers (Beckton Dickinson), or sorted on a FACSAria. Data were analyzed using FlowJo Version 9.2 software (TreeStar) with populations defined by size, viability, and “fluorescence minus one” isotype controls.

### Immunofluorescence

Alexa-CCL22 labelled ACKR2+ fibroblasts were flow-sorted from a single cell suspension using the ARIAII (Beckman and Dickinson) as per the flow protocol. ACKR2^+^ Fibroblasts were resuspended in PBS at a density of 2000 cells per ml. 100µl of this cell suspension was loaded into a cytospin 3 machine (Thermo-Shandon) and the cells were spun for 5 minutes at 200rpm onto Superfrost™ plus slides. The cells were air dried in the dark and mounted in Vectashield Hard set mounting medium (Vector Laboratories). The cells were visualised using a Zeiss Spinning Disc confocal microscope.

### Fibroblast culture and analyses

Primary ACKR2 positive fibroblasts were flow-sorted from a single cell suspension using the ARIAII (Beckman and Dickinson) as per the flow protocol. Retrieved cells were spun down at 400g for 5 minutes and washed 3 times into EMEM with 15% FBS, 1X Penicillin/Streptomycin, non essential amino acids, and sodium pyruvate. The cells were plated at a density of approximately 5×10^3^ cells per well of a 24 well plate (Gibco). Fibroblasts were cultured as standard in an incubator at 37°C, 5% CO_2_.

### Statistics

Statistical tests were carried out using Graph Pad Prism software and the individual tests used are indicated in the relevant figure legends. P=0.05 was taken as a cut-off for statistical significance.

## Results

### ACKR2 is stromally expressed in the lung

Data on ACKR2 expression profiles available through the Immgen database (www.immgen.org) reveal that the lung is the tissue with the highest expression (Supplemental Figure 1). We have previously described the versatile use of fluorescently labelled chemokines, instead of antibodies, to detect their cognate receptors using flow cytometry as a read out(35–38). We used this approach with Alexa-647 labelled CCL22 (a high affinity ACKR2 ligand: Alexa-CCL22) uptake to detect ACKR2 in lung digests. Flow cytometric analysis of Alexa-CCL22 stained lung digests failed to detect any significant expression on CD45+ leukocytes in either WT or ACKR2-/-lungs (Figure 1a). This therefore indicated that ACKR2 expression in the lung was predominantly stromal in origin. We used RNA sequencing to generate data on the transcriptional profile of FACS-sorted pulmonary stromal cell types at rest and over the course of an influenza infection to examine possible pathogen-driven alterations in expression. As shown in Figure 1b, ACKR2 expression was essentially undetectable in epithelial cells but was present at low levels in fibroblasts and at very high levels in blood endothelial cells. Expression did not vary significantly over the course of influenza infection. Examination of pulmonary ACKR2 expression by qPCR from embryonic day 13.5 to 9 weeks of age indicated that expression is low within the embryo but that it is markedly upregulated immediately after birth and presumably coincident with the onset of breathing. This increased level is maintained and increased as mice age (Figure 1c).

**Figure 1.**
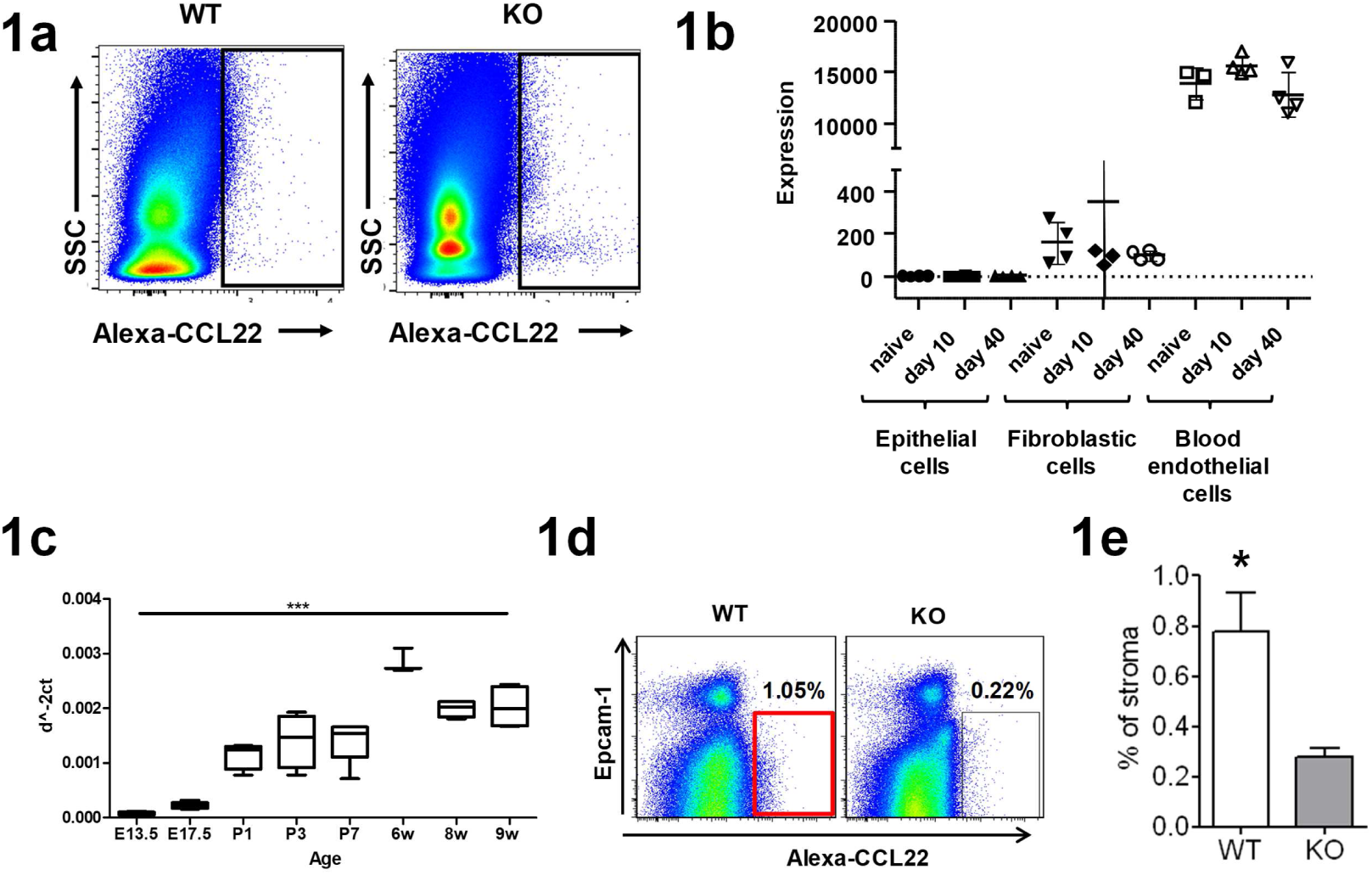
In vitro analysis of pulmonary ACKR2 expression. A) flow cytometric analysis showing lack of expression of ACKR2 by CD45+ve cells in lung digests as assessed by comparing Alexa-CCL22 uptake (X-axis) in WT and ACKR2-/- (KO) lungs. The Y axis shows side scatter (SSC). B) analysis of transcriptomic data generated from lung stromal populations showing low level ACKR2 expression in fibroblasts but high-level expression in blood endothelial cells. These data were generated from resting lung stromal populations as well as from the same populations retrieved from lungs at the indicated times (10 and 40 days) post influenza virus infection. C) qPCR analysis of pulmonary ACKR2 expression from embryonic day E13.5 to adult 9 week old mice. Data were normalised to expression of the housekeeping gene Tata-binding protein. Statistical significance was tested using a 1-way ANOVA test and ***=p<0.001. D) flow cytometric analysis of CD45-ve Alexa-CCL22 internalising cells from WT and ACKR2-/- lungs. The X-axis shows the Alexa-CXCL22 binding whilst the Y-axis shows staining for the epithelial marker EpCAM. E) quantification of the percentage of ACKR2+ve stromal cells detected using this Alexa-CCL22 in vitro labelling approach. Statistical significance was tested using Student’s T test and *=p<0.05.

Next, we tried to use the in vitro Alexa-CCL22 detection method to examine ACKR2 expression on non-leukocytic stromal cells in the digested lung. However, and as shown in Figure 1d, this technology, which works well with leukocytes(35, 38, 39) and mammary gland fibroblasts(30), revealed only a minor difference in the numbers of Alexa-CCL22 internalising cells in WT or ACKR2-/- lungs. These data are summarised numerically in Figure 1e.

Overall, therefore, these data indicate that ACKR2 is predominantly expressed on stromal cells within the lung but that flow cytometry utilising fluorescent chemokine uptake with digested lung tissue has limited sensitivity to detect the key stromal expressing cell types.

### Intra-tracheal fluorescent chemokine administration identifies key ACKR2-expressing stromal components

We reasoned that the function of stromal ACKR2 expression may be dependent on interactions with other stromal components and that the inability to detect it using flow cytometry reflects the inability to take up Alexa-CCL22 due to absence of these interactions. We therefore harvested intact lungs, inflated them with Alexa-CCL22 (in RPMI) intra-tracheally followed by incubation at 37°C for 1 hour (Figure 2a). Following this, flow cytometric analysis of digested lungs focusing on CD45-ve cellular populations now revealed a sizeable population of Alexa-CCL22 internalising cells in WT lungs which are absent from ACKR2-/- lungs (Figure 2b). These cells are non-epithelial as they are negative for EpCAM staining and these data are summarised numerically in Figure 2c. Further flow cytometric analysis indicated that these cells fall into 3 basic categories as defined by CD31 and Gp38 expression (Figure 2d). These represent fibroblasts (R1; CD31-Gp38-), lymphatic endothelial cells (R2; CD31+Gp38+) and blood endothelial cells (R3; CD31+Gp38-) which are enumerated as shown in Figure 2e.

**Figure 2.**
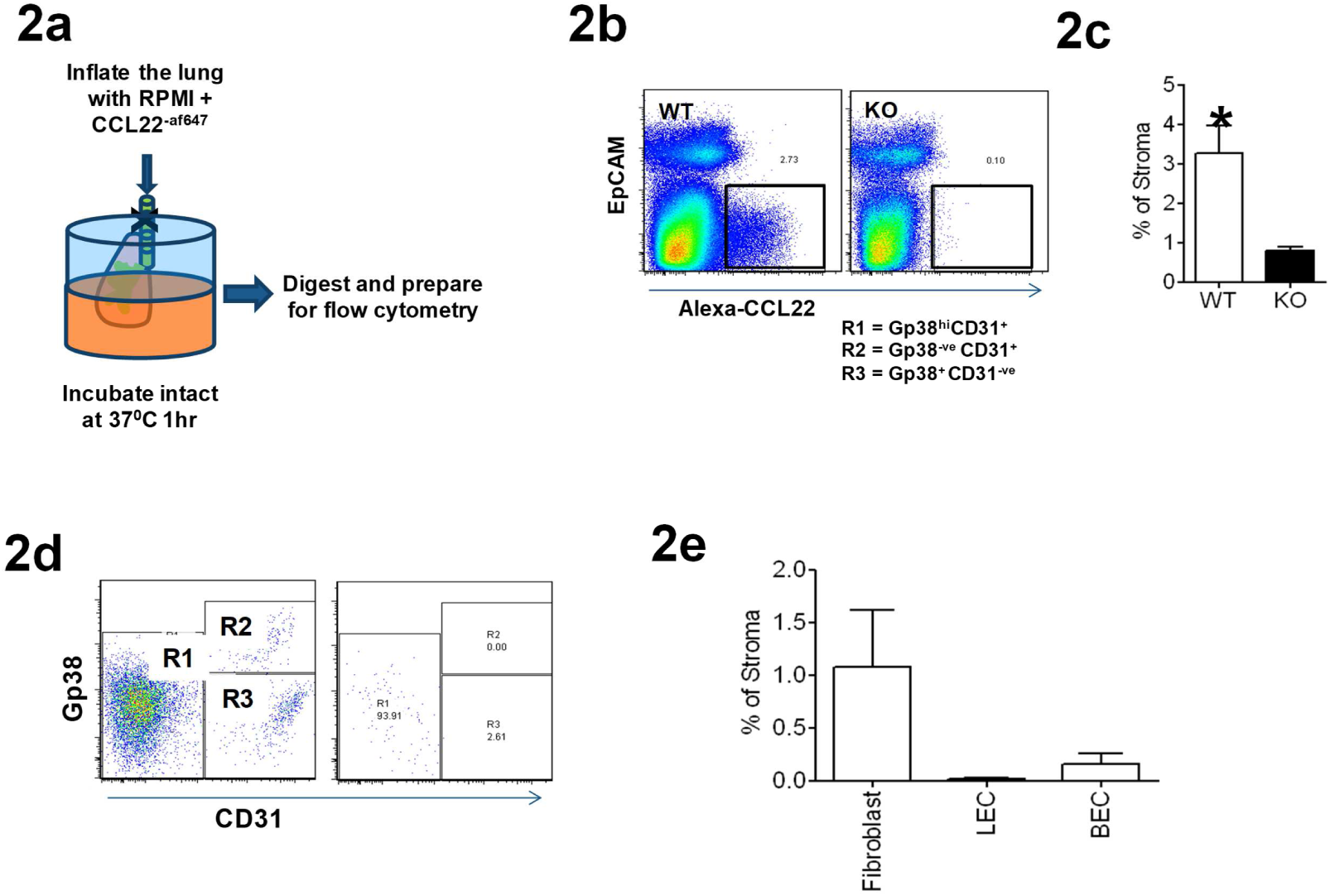
Detection of stromal ACKR2 expression by intra-tracheal administration of Alexa-CCL22. A) diagram indicating the basic methodology which involves removing intact lungs and inflating them through the trachea using an RPMI/Alexa-CCL22 mix. The lung is then incubated at 37°C for 1 hour then digested and prepared for flow cytometry. B) flow cytometric analysis showing the presence of a significant Alexa-CCL22 internalising stromal component in the lung. The X-axis shows the Alexa-CXCL22 binding whilst the Y-axis shows staining for the epithelial marker EpCAM. C) quantification of the levels of stromal ACKR2 detected in WT and ACKR2-/- (KO) lungs following intra-tracheal Alexa-CCL22 administration. Statistical significance was tested using Student’s T test and *=p<0.05. D) flow cytometric analysis of the CD45-ve EpCAM-ve non-epithelial stromal component expressing ACKR2 on the basis of CD31 and Gp38 expression. Data shown are from Alexa-CCL22 binding lung stromal cells from WT (left plot) and ACKR2-/- (right plot) mice. E) summary analysis of the percentage of CD45-ve stroma associated with each of the indicated cell types following intratracheal administration of Alexa-CCL22.

Overall therefore these data demonstrate that it is possible to detect stromal cell populations that bind and internalise Alexa-CCL22 via ACKR2 by introducing the chemokine intra-tracheally into the intact lung. They also demonstrate novel stromal expression patterns for ACKR2 within the lung.

### A subpopulation of Fibroblasts in the lung express ACKR2

Cells identified in the R1 gate in Figure 2d, which were negative for markers of lymphatic and vascular endothelial cells, were further phenotyped. Initially these cells were isolated by cell-sorting and then grown in tissue culture. As shown in Figure 3a, the cells display a morphology suggestive of a fibroblastic phenotype. Figure 3b further shows that the ACKR2+ fibroblastic cells were capable of binding and internalising Alexa-CCL22 (arrow) confirming expression of functional ACKR2 (Figure 3b). Further flow cytometric analysis revealed that these cells are negative for markers of epithelial (CD166) and endothelial (CD49f) cells but positive for the fibroblastic marker CD140a (Figure 3c). Overall these data indicate that one of the ACKR2-expressing stromal components in the lung detected by intra-tracheal Alexa-CCL22 administration is a fibroblastic subpopulation characterised by CD140a expression.

**Figure 3.**
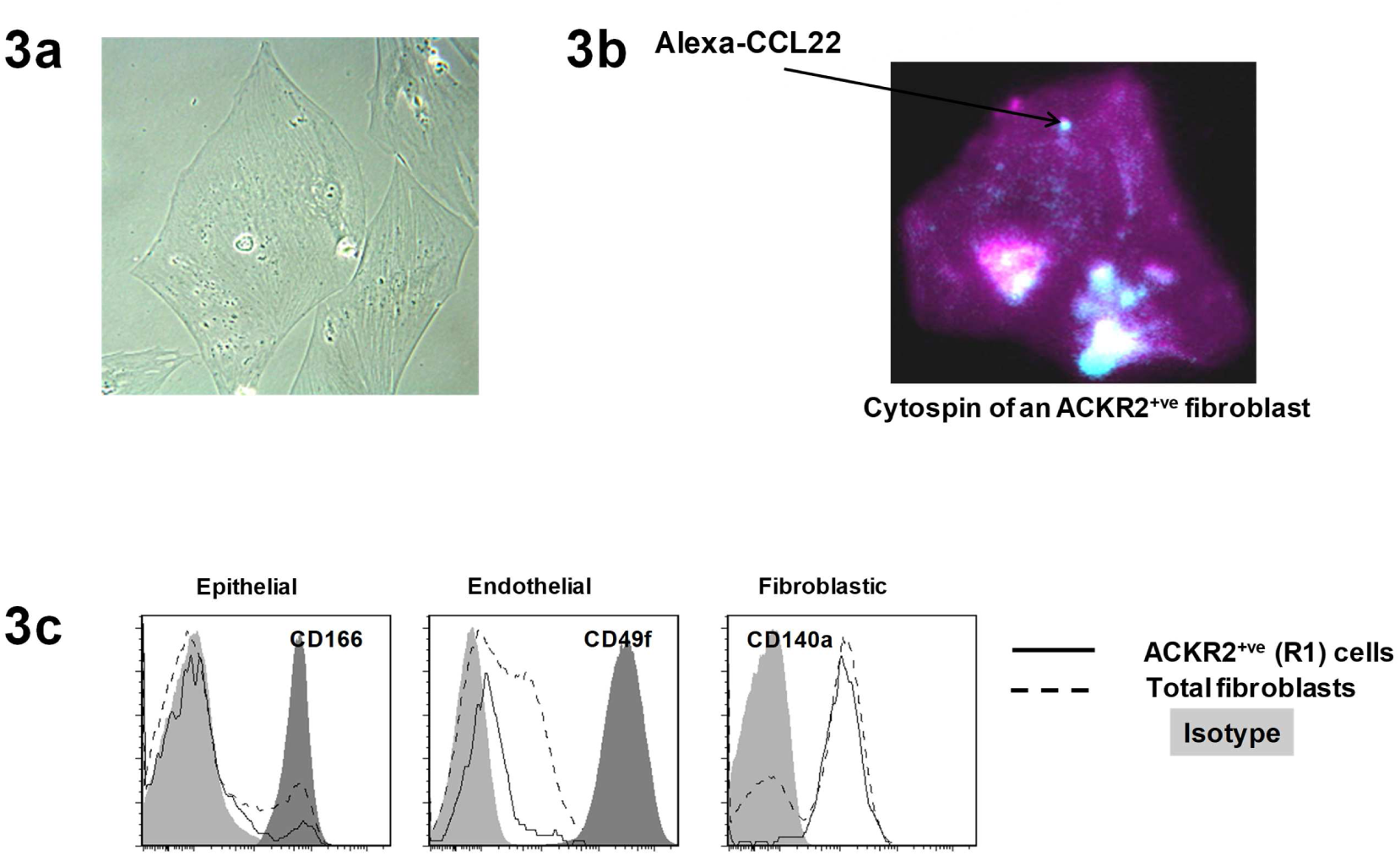
Identification of a fibroblastic population as a component of the ACKR2-expressing stromal population. A) brightfield image of cells sorted for internalisation of fluorescently labelled Alexa-CCL22. These cells are negative for Gp38, CD45 and CD31 expression. B) fluorescent confocal image of ACKR2+ve fibroblastic cells binding and internalising Alexa-CCL22 (arrowed). C) flow cytometric analysis of the fibroblastic population showing absence of expression of epithelial (CD166) and endothelial (CD49f) markers but positive expression of fibroblastic markers (CD140a).

### Intravenous fluorescent chemokine administration identifies additional ACKR2-expressing stromal components

We have previously reported expression of ACKR2 in lymphatic endothelial cells(31), however expression in blood endothelial cells as shown in Figures 2d and 2e has not been reported. In fact, and as shown in Supplemental Figure 2, comparison of transcriptomic data from a broad range of microvascular endothelial types indicates that expression in blood endothelial cells is peculiar to the lung. Given that we have reported polarisation of expression of ACKR2 in trophoblasts and lymphatic endothelial cells(31, 34, 40) we wondered whether it may also be polarised in blood endothelial cells. If this is the case then it is possible that the vascular facing aspect of endothelial cells may be more able to internalise Alexa-CCL22 and thus the blood endothelial cell component might be underestimated by the intra-tracheal administration of the chemokine. We therefore next injected Alexa-CCL22 intravenously (Figure 4a). Again, flow cytometric analysis revealed Alexa-CCL22 uptake by 3 populations of non-epithelial stromal cells i.e. fibroblasts, blood endothelial cells and lymphatic endothelial cells (Figure 4b). However, and in contrast to intra-tracheal administration, intravenous administration detected blood endothelial cells as being by far the dominant ACKR2-expressing stromal population. Figure 4c shows that intravenous administration highlights blood endothelial cells as being the predominant stromal cell component in the lung and comparing intra-tracheal administration with intravenous administration (Figure 4d) reveals the differences in cellular detection using these two approaches. As we have previously shown expression of ACKR2 by lymphatic endothelial cells(31, 32) it remained possible that what we have characterised as lung blood endothelial cells are in fact lung lymphatic endothelial cells displaying an altered CD31/Gp38 phenotype compared to other lymphatic endothelial cell populations. To formally test this we examined lung lymphatic endothelial cell expression of CD31 and Gp38 using Prox-1 reporter mice. Prox-1 is a definitive marker and an essential master regulator, of lymphatic endothelial cells(41). As shown in Figure 4e, Prox-1+ve cells from reporter mouse lungs were exclusively co-positive for CD31 and Gp38 confirming the faithfulness of the lymphatic phenotype in the lung. These data further confirm the blood endothelial nature of the ACKR2+ve stromal cells.

**Figure 4.**
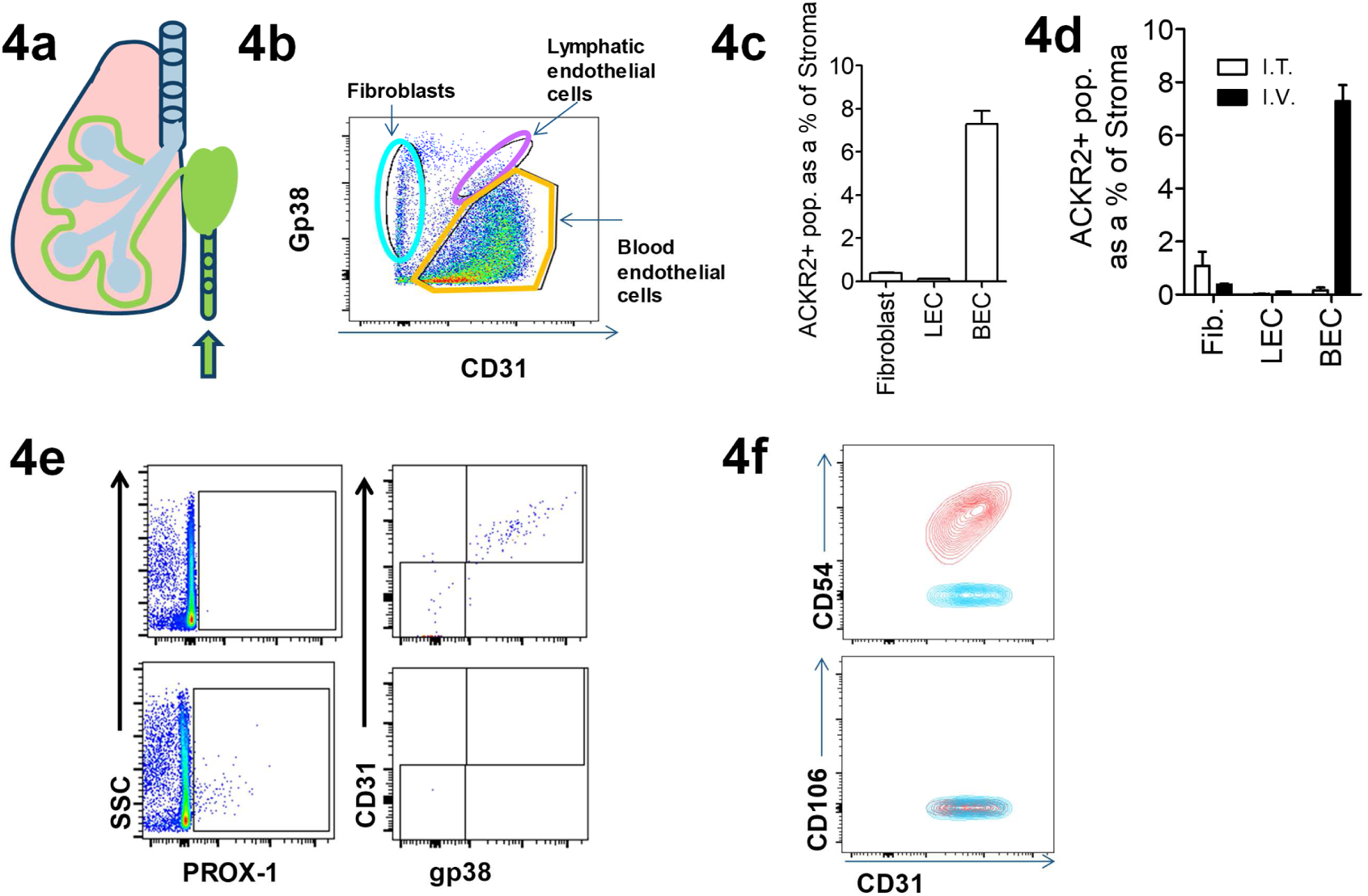
Intravenous administration of Alexa-CCL22 reveals strong expression of ACKR2 on the luminal face of pulmonary vascular endothelial cells. A) diagram indicating intravenous injection of Alexa-CCL22 into live mice. B) flow cytometric analysis of CD31 (X-axis) and Gp38 (Y-axis) expression by CD45-ve EpCAM-ve stromal cells that internalise Alexa-CCL22. The identified populations are indicated. C) quantification of the % of CD45-ve stromal cellular populations internalising Alexa-CCL22 following intravenous administration of Alexa-CCL22. D) comparison of intratracheal and intravenous administration and the impact on the relative size of the ACKR2+ stromal cell populations detected. Fib.-fibroblasts; LEC-lymphatic endothelial cells; BEC-blood endothelial cells. E) flow cytometric analysis of the CD31+ve, Alexa-CCL22 internalising blood endothelial cells indicate that they are positive for expression of CD54 and negative for CD106 and thus are alveolar, rather than peribronchial, endothelial cells. Red-the gated CD31+ cells that have been antibody stained; Blue-fluorescence minus one (FMO) control.

In the lung there are two major blood vascular beds: one associated with bronchial tissues and one with alveolar tissues. These can be discriminated on the basis of CD54 and CD31 expression(42). As shown in Figure 4f the Alexa-CCL22 internalising blood endothelial cell population is strongly co-positive for CD31 and CD54 demonstrating that this population is associated with alveolar blood vessels and not peribronchial blood vessels.

### In situ hybridisation and antibody expression confirm blood endothelial cell expression of ACKR2

To further validate ACKR2 expression by murine pulmonary vascular endothelial cells, we carried out in situ hybridisation. As shown in Figure 5a, clear in situ hybridisation signals were seen associated with alveolar endothelial cells in blood vessels surrounding the alveolar air space in WT, but not ACKR2-/-, adult murine lungs. A higher magnification image of a portion of Figure 5ai is shown in Figure 5b further highlighting the peri-alveolar localisation of the vascular staining. Therefore, in situ hybridisation confirms blood endothelial cell expression of ACKR2 in the murine lung.

**Figure 5.**
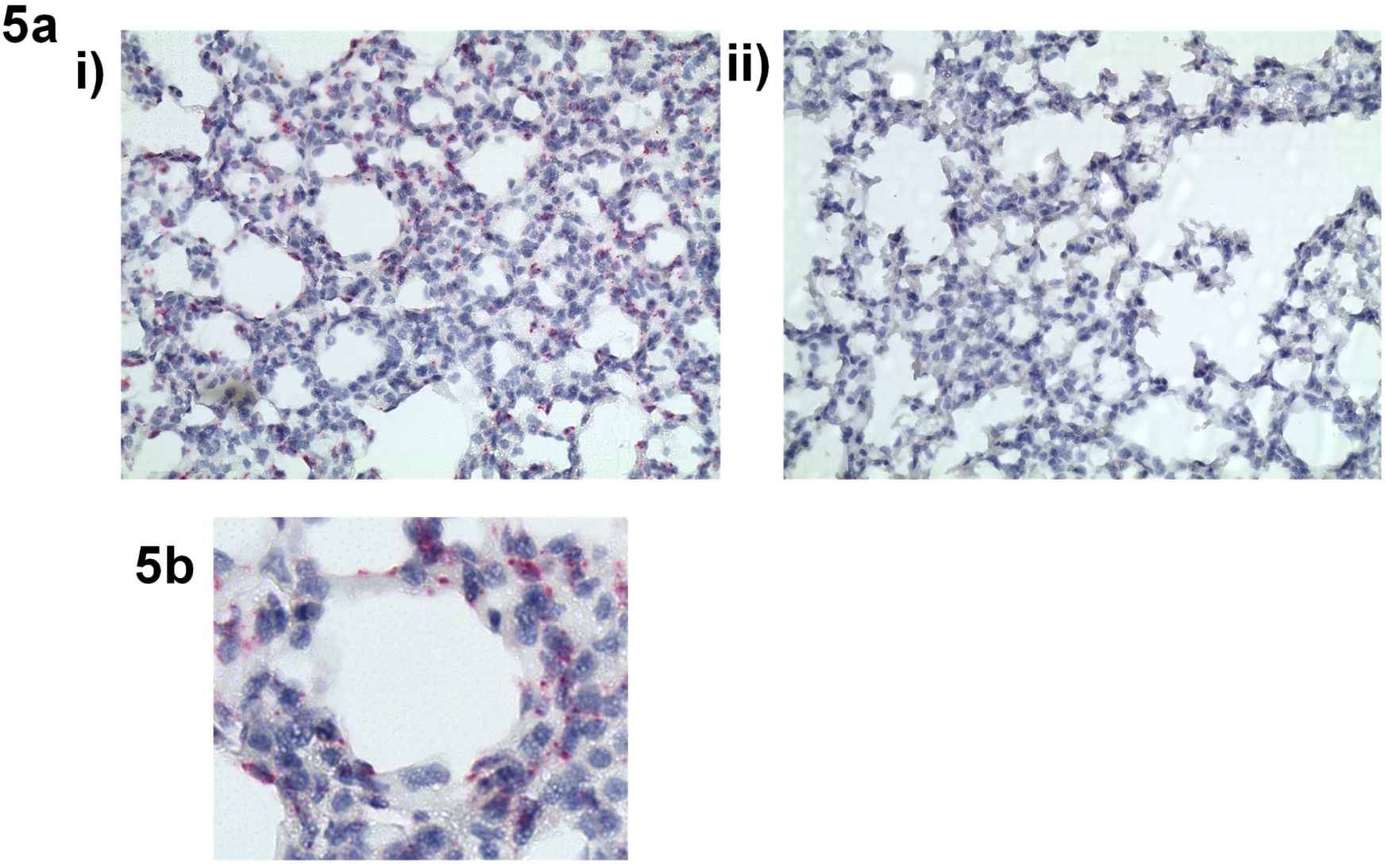
in situ and antibody detection of ACKR2 on pulmonary alveolar blood endothelial cells in mouse and human lung. A) in situ hybridisation showing positivity for ACKR2 expression in alveolar endothelial cells in murine lungs (arrow). i) WT lungs and ii) ACKR2-/- lungs. Note the absence of signal in alveolar macrophages. B) high power image cropped from Figure 5i showing the peri-alveolar localisation of the vascular staining.

## Discussion

Atypical chemokine receptors are predominantly expressed on stromal cell types and serve key functions in localising and fine-tuning chemokine activities(15). Whilst a number of antibodies and reporter mouse-based approaches are available for analysis of atypical chemokine receptor expression patterns, in many cases these are limited and applicable only to mouse and humans. Given that many of the atypical chemokine receptors display strong evolutionary conservation, other more versatile approaches would therefore represent a significant improvement in the methodological repertoire for atypical chemokine receptor expression analysis. Here we demonstrate, using the lung, that fluorescently-labelled chemokines can be used, with intact tissues, to precisely isolate and phenotypically define stromal cell types expressing individual atypical chemokine receptors. This is particularly important when, as in the current analyses, removal of the cells from their stromal environment impairs ACKR2 function and thus frustrates this detection methodology. The ability to chemically synthesise chemokines with relative ease and to introduce discrete fluorescent markers at the carboxy terminus(39) means that this approach has full versatility and is appropriate for all members of the chemokine receptor family and all species expressing either typical or atypical chemokine receptors. Importantly, whilst we demonstrate the utility of this approach using WT and KO mice, the use of appropriate unlabelled competing chemokines specific for the receptor being studied will allow this technology to be used under circumstances, or in species, where KO models do not exist.

Here we show that intra-tracheal, and intra-venous, administration of fluorescently labelled CCL22 is capable of identifying stromal cell populations expressing ACKR2. Importantly, in the context of vascular endothelial cell expression, intravenous administration has the advantage of detecting chemokine receptor expression with a polarity favouring expression on the luminal side of the endothelium. Together these approaches allowed us to define fibroblasts, lymphatic endothelial cells, and surprisingly alveolar blood endothelial cells as key sites of stromal ACKR2 expression in the lung. Surprisingly, and in contrast to previous reports, we did not detect ACKR2 activity on alveolar macrophages(43, 44). This may be due to species differences in expression or alternatively may be a consequence of non-specific antibody internalisation by the alveolar macrophages. Further analysis is required to address this discrepancy.

Importantly this technology is applicable to other atypical chemokine receptors as well as to the classical chemokine receptors and to any tissue which can be incubated in vitro with fluorescent chemokines. This approach would, for example, be readily amenable for use with intestinal tissue, liver and skin.

In summary therefore we report a methodology appropriate for detecting atypical chemokine receptor expression in the lung which we believe to be sufficiently versatile to be useful to detect other atypical and typical chemokine receptors in numerous tissues in vivo in divergent species. The data indicate for the first time the stromal expression patterns of ACKR2 in the lung with notably high expression levels on alveolar endothelial cells. These data will help interpret the outcome of analysis of ACKR2 function the lung which remains poorly defined.

