## Supplementary Information for "A novel methodology for defining stromal expression of atypical chemokine receptors in vivo"

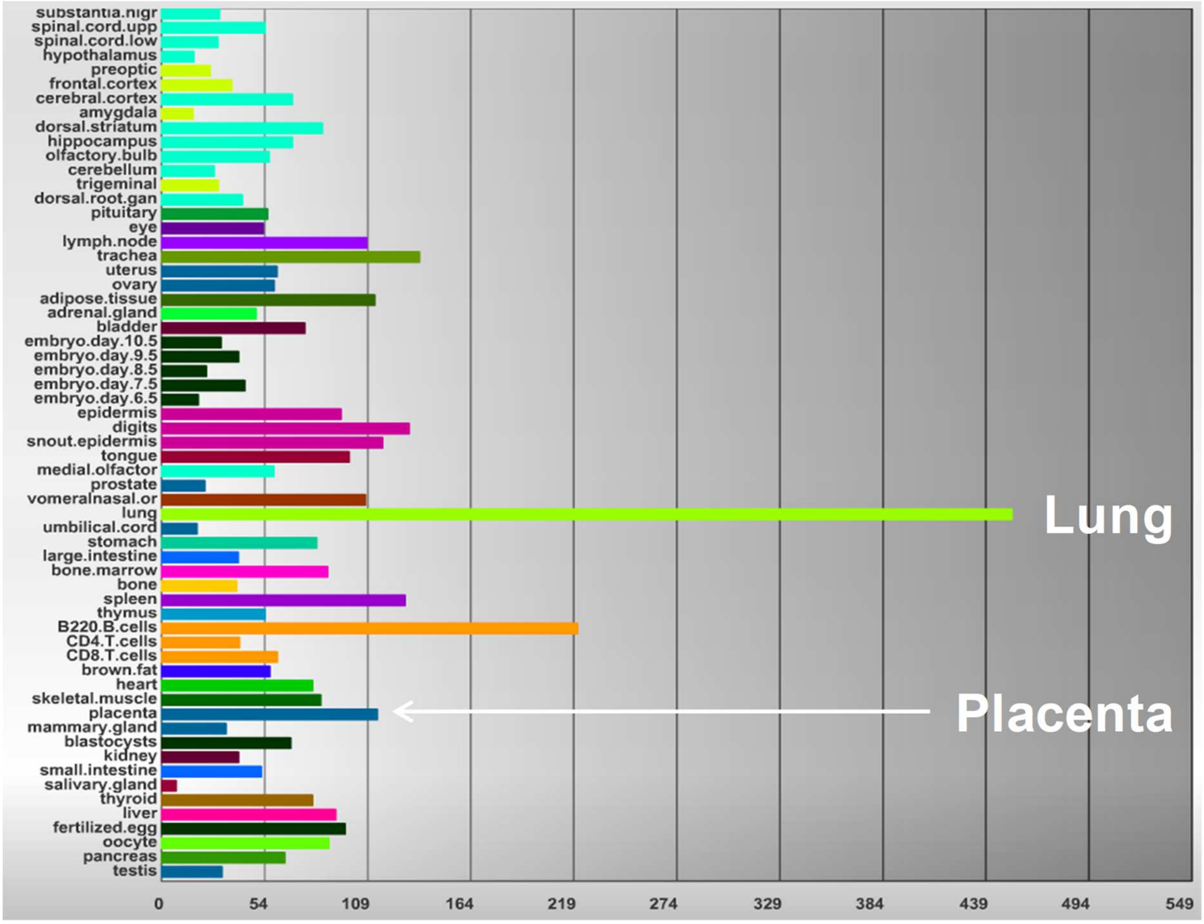


**Supplemental Figure 1:** summary diagram showing expression analysis from Immgen indicating the strong expression of ACKR2 in mouse lung.

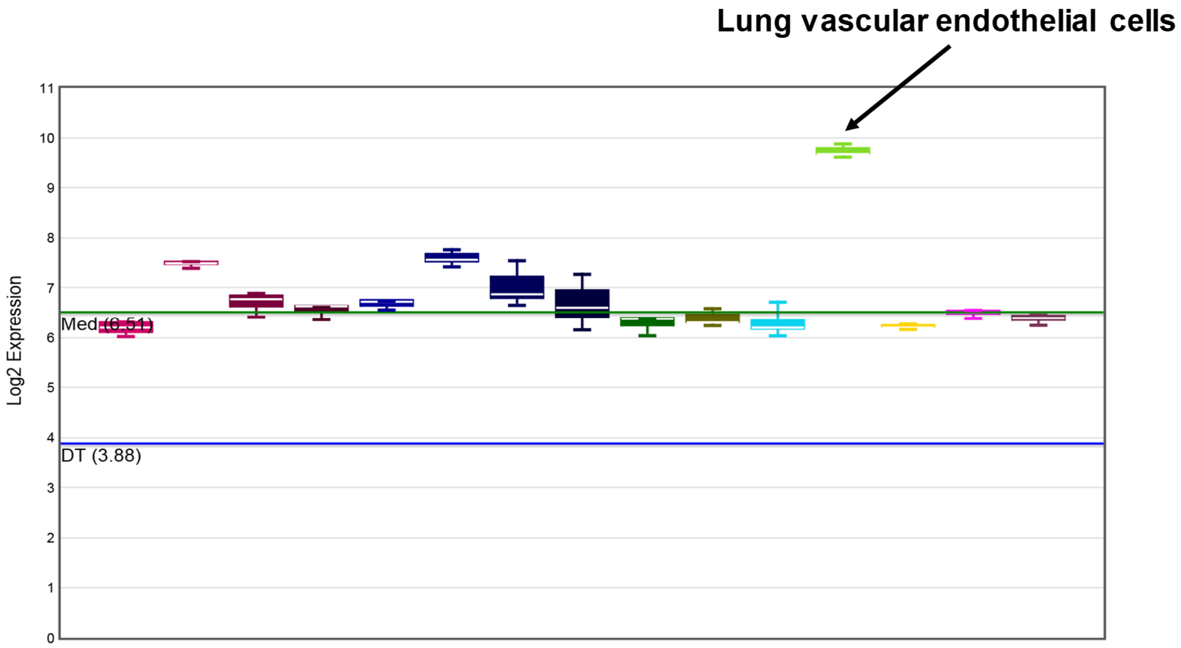


**Supplemental Figure 2:** in silico analysis of the molecular signatures of tissue-specific microvascular endothelial cells for ACKR2 expression. Lung endothelial cells expression is indicated by the arrow.
